# Neuropathology of RAN translation proteins in Fragile X-associated Tremor/Ataxia Syndrome

**DOI:** 10.1101/672444

**Authors:** Amy Krans, Geena Skariah, Yuan Zhang, Bryana Bayly, Peter K. Todd

## Abstract

CGG repeat expansions in *FMR1* cause the neurodegenerative disorder Fragile X-associated Tremor/Ataxia Syndrome (FXTAS). Ubiquitinated neuronal intranuclear inclusions (NIIs) are the neuropathological hallmark of FXTAS. Both sense strand derived CGG repeats and antisense strand derived CCG repeats support non-AUG initiated (RAN) translation of homopolymeric proteins in potentially 6 different reading frames. However, the relative abundance of these proteins in FXTAS brains and their co-localization with each other and NIIs is lacking. Here we describe rater-blinded assessment of immunohistochemical and immunofluorescence staining with newly generated antibodies to different CGG RAN translation products in FXTAS and control brains as well as co-staining with ubiquitin, p62/SQSTM1, and ubiquilin 2. We find that both FMRpolyG and a second CGG repeat derived RAN translation product, FMRpolyA, accumulate in aggregates in FXTAS brains. FMRpolyG is a near-obligate component of both ubiquitin-positive and p62-positive NIIs in FXTAS, with occurrence of aggregates in 20% of all hippocampal neurons and >90% of all inclusions. A subset of these inclusions also stain positive for the ALS/FTD associated protein ubiquilin 2. Ubiquitinated inclusions and FMRpolyG+ aggregates are rarer in cortex and cerebellum. Intriguingly, FMRpolyG staining is also visible in control neuronal nuclei. In contrast to FMRpolyG, staining for FMRpolyA and CCG antisense derived RAN translation products were less abundant and were infrequent components of FMRpolyG+ inclusions. In conclusion, RAN translated FMRpolyG is a common component of ubiquitin and p62 positive inclusions in FXTAS patient brains.

## Introduction

Nucleotide repeat expansions cause more than forty neurological diseases, including Huntington disease, C9ORF72-associated Amyotrophic Lateral Sclerosis and Frontotemporal Dementia, and a number of spinocerebellar ataxias and myotonic dystrophies [20]. Repeat expansions cause disease through a series of overlapping mechanisms, including loss of expression of the repeat containing gene, gain of function toxicity elicited by repeat RNA, and gain of function toxicity elicited by proteins generated from AUG initiated or repeat associated non-AUG initiated (RAN) translation [17, 18].

Fragile X-associated Tremor/Ataxia Syndrome (FXTAS) is a neurodegenerative disease caused by a trinucleotide CGG repeat expansion in the 5’ untranslated region (UTR) of the *FMR1* gene [8]. Clinically, FXTAS is characterized by intention tremor, ataxia, gait abnormalities and cognitive decline [2]. Both patients and CGG knock-in (KI) mouse models of disease have elevated *FMR1* mRNA but lower basal expression of the protein product, FMRP [26, 27]. The pathologic hallmark of FXTAS is the accumulation of ubiquitinated neuronal intranuclear inclusions (NIIs) throughout the brain [7, 25]. NIIs are most prominent in the hippocampus and, to a lesser degree, in the frontal cortex and granule cell layer of the cerebellum [6]. Astrocytic inclusions also occur frequently within the brainstem and other brain regions [6, 7]. Despite their clear role in the clinical syndrome and evidence of cerebellar atrophy on both pathological analysis and imaging, ubiquitinated inclusions are relatively rare in cerebellar Purkinje neurons [1].

In initial work in FXTAS, no single dominant protein species was found in these aggregates. Proteins identified by mass spectrometry and immunohistochemically include, but are not limited to ubiquitinated proteins, lamin A/C, αB crystallin, a series of histone proteins and proteasomal subunits, and the RNA binding proteins Sam68, Muscleblind1, and hnRNPA2/B1 [12, 13, 23, 24]. In addition, biotinylated antisense RNA probes targeting the 5’ UTR, coding region and 3’UTRs of *FMR1* diffusely stained inclusions in nuclei isolated from FXTAS patient cortex [28]. Based on these initial findings, it was proposed that CGG repeat RNA serves as a nidus for inclusion formation by binding to and sequestering specific proteins into these aggregates. Consistent with this model, many of the factors identified within inclusions to date are RNA binding proteins that associate with CGG repeat RNA in *in vitro* assays [13, 22–24]. Of note, FMRP itself is not found in the inclusions and loss of the protein is not associated with neurodegeneration in clinical cases or animal models [11, 14].

An alternative mechanism by which inclusions may form in FXTAS is based on a unique form of protein translational initiation known as Repeat Associated non-AUG (RAN) translation [5, 18]. In rabbit reticulocyte lysates, transfected cells and neurons, Drosophila, and mouse models of the disease, expression of the 5’ UTR of *FMR1* leads to RAN translation of a series of homopolymeric proteins, with different efficiencies of production and accumulation in different reading frames [15, 19, 21, 29]. RAN translation can occur from both sense strand CGG repeat (producing polyglycine (FMRpolyG), polyalanine (FMRpolyA) and polyarginine (FMRpolyR) repeat containing proteins) and antisense strand CCG repeat (producing polyproline (ASFMRpolyP), polyalanine (ASFMRpolyA), and polyarginine (ASFMRpolyR) containing proteins) mRNA transcripts in reporter assays [15, 16]. FMRpolyG production in particular appears critical for NII formation, as mutations that largely preclude FMRpolyG production in the sequence just 5’ to the repeat prevents NII formation in *Drosophila*, and both CGG KI mice and repeat expressing transgenic mice [10, 19, 21, 29]. Moreover, generation of FMRPolyG absent the CGG repeat through use of alternative codons is sufficient to elicit inclusion formation in transfected cells [9]. Importantly, the ability to generate FMRpolyG is predictive of whether CGG repeats expressed in neurons, *Drosophila*, or in transgenic mice are capable of eliciting neurodegeneration despite comparable levels of expression of the repeat containing mRNA [21, 29].

Previous studies have demonstrated the presence of FMRpolyG positive aggregates in FXTAS patient tissue using different mouse monoclonal antibodies targeted against either the N-terminal region or C-terminal region of FMRpolyG [3, 21, 29]. In addition, ASFMRpolyP and ASFMRpolyA staining is detected in some aggregates in FXTAS cases [16]. However, the relative abundance of each RAN peptide in FXTAS has not been systematically evaluated to date. Moreover, direct comparisons of RAN pathology with ubiquitinated inclusion burden have not yet been performed in any systematic fashion.

Here we describe a series of new antibodies generated against FMRpolyG and FMRpolyA epitopes that contain a portion of the homopolymeric repeat. After thoroughly establishing the specificity of these antibodies for their pathological targets, we use them to determine the relative distribution and abundance of different RAN derived proteins in the brains of FXTAS patients and their overlap with ubiquitin and p62 pathology in a rater-blinded fashion to provide unbiased and rigorous results. These studies suggest that FMRpolyG is a near-obligate component of NIIs in FXTAS and further support a role for FMRpolyG accumulation in the FXTAS pathology.

## Materials and methods

### FMR polyclonal antibody generation

Rabbit polyclonal antibodies were generated by Abclonal (Cambridge, MA). Peptides were generated corresponded to either the N-terminal or C-terminal region of the predicted proteins and extended into the repeat region. Exact sequences can be found in **Figure 2**. Antibodies were affinity purified from anti-sera. In addition to receipt of the antibodies, pre- and post-bleed serum samples from each rabbits and the peptides used to generate the antibodies were also received.

Pre-bleed sera and the peptide were used to characterize the antibodies. Protein concentrations of the pre-bleed sera were determined. The dilution of pre-bleed sera used had an equal protein concentration as the dilution of antibody used. A 100x excess of peptide was incubated with the antibody prior to use to block the antibodies’ ability to bind to antigen.

### Western blotting

HEK293 cells were transfected with constructs expressing FMRpolyG, FMRpolyA, or EGFP-N1 using Fugene HD (Promega) following the manufacturer’s protocol. Lysates were collected 24 hours after transfection, as previously described [3, 21, 29]. Proteins were separated using SDS-PAGE and transferred to PVDF membrane. Proteins were detected with antibodies against GFP (Sigma, 1:1000) and FMR polyclonal antibodies (1:1000). GAPDH or tubulin was used as a loading control.

### Immunocytochemistry

HEK293 cells were grown on a 4-well chamber slide and transfected with FMRpolyG, FMRpolyA, or EGFP-N1 using Fugene HD. Cells were fixed with 4% paraformaldehyde 24 hours after transfection and permeabilized with 0.1% Triton X-100 in phosphate buffered saline, 10mM MgCl_2_, 1mM CaCl_2_ (PBS-MC). 5% normal goat serum in PBS-MC, 0.1% Triton was used as a blocking agent and diluent for primary antibodies. N-terminal FMRpolyG (NTF1, 1:200), C-terminal FMRpolyG, (CTF1, 1:100), and FMRpolyA (1:100) were incubated with GFP (1:1000) overnight at 4°C. AlexaFluor488 goat anti-mouse and AlexaFluor555 goat anti-rabbit IgG secondary antibodies were used. Nuclei were stained and coverslips mounted using Prolong Gold with DAPI (ThermoFisher). Images were captured on an inverted Olympus (Tokyo, Japan) IX71 microscope at the same exposure and processed using SlideBook 5.5 software.

### Immunohistochemistry and co-immunofluorescence

Control and FXTAS brain regions were obtained from the University of Michigan Brain Bank, the New York Brain Bank, and the University of Florida. FXTAS cases 1-3 were described previously [16, 29].

For immunohistochemistry, paraffin embedded sections were deparaffinized with a series of xylene washes and decreasing concentrations of ethanol. Antigen retrieval (AR) was done with citrate buffer, if necessary. Endogenous peroxidase was quenched with 1% hydrogen peroxide for 30 minutes. Sections were blocked in 5% normal goat serum (NGS) in Tris, pH7.6. Primary antibodies, p62 (Proteintech, 1:1000, acid AR), ubiquitin (DAKO, 1:250, basic AR), ubiquilin 2 (Novus Biologicals, 1:200, acid AR), NTF1 (Abclonal, 1:200, no AR), CTF1 (Abclonal, 1:40, basic AR), and FMRpolyA (Abclonal, 1:100, basic AR) were diluted in 5% NGS, Tris pH7.6, 0.1% Triton-X 100, 0.5% bovine serum albumin (Tris B) and were incubated with sections overnight at 4°C. The following day, sections were washed in Tris pH7.6, 0.1% Triton (Tris A) and Tris B. Antibody detection was determined using VECTASTAIN Elite ABC HRP kit (Vector Laboratories) following the manufacturer’s protocol. Sections were counterstained with hematoxylin (Vector Laboratories). After dehydrating the slides through a series of increasing concentration of ethanol and then xylenes, coverslips were mounted to the slides using DPX mounting media. Images were taken on an Olympus BX51 microscope.

For coimmunofluorescence, sections were blocked with 5% NGS in PBS-MC. Ubiquitin, ubiquilin 2, p62, NTF1, CTF1, and FMRpolyA were diluted in 5% NGS-PBS-MC, overnight at 4°C. Goat anti-rabbit and goat anti-mouse IgG antibodies conjugated to AlexaFluor488 or AlexaFluor635 (ThermoFisher, 1:500) were used. Nuclei were stained and coverslips mounted using Prolong Gold with DAPI (ThermoFisher). Images were taken on an Olympus confocal microscope and compiled and analyzed using ImageJ.

### Analysis of immunohistochemistry

Intensity of staining was graded on a four-point integral scale of 0 (no staining) to 3 (intense staining). An additional point was added if an aggregate/inclusion was present. Grading was done by three separate reviewers blinded to the genotype of the tissue as well as the identity of the antibody being used. All reviewers were trained and tested to ensure consistent grading for intensities and inclusion identification. A subset of images was repeated among reviewers to assess variability during retest. All images were randomized and anonymized prior to assigning them to the reviewers. An average of 20 images per slide was counted for each of the control and FXTAS cases and multiple brain regions were analyzed.

### Statistical analysis

Tabulated scores of staining intensity and percentage of cells containing an aggregate were analyzed using GraphPad Prism. As this was a non-parametric scale, a Mann-Whitney U two-tailed t-test was used to determine if there were differences in the number of aggregates per neuron.

## Results

Ubiquitin, p62/SQSTM1, and ubiquilin 2 (UBQLN2) are components of intracellular inclusions in a number of neurodegenerative diseases, including Huntington disease and C9ORF72-associated ALS/FTD. In non-disease brain, these proteins are generally diffuse within neurons when visualized by immunohistochemistry in both the hippocampus and cortex (Fig. 1A, 1D and 1G). Consistent with the original pathological description of FXTAS, we observed significant ubiquitin pathology in FXTAS cases compared to controls (Fig. 1A), with intranuclear inclusions observed throughout the hippocampus and cortex, but with relatively fewer inclusions in the brainstem and cerebellum. Intranuclear neuronal aggregates in FXTAS also stained positive for p62 (Fig. 1D), consistent with a recent report [4]. We also observed UBQLN2 positive inclusions in FXTAS neuronal nuclei (Fig. 1G).

**Figure 1.**
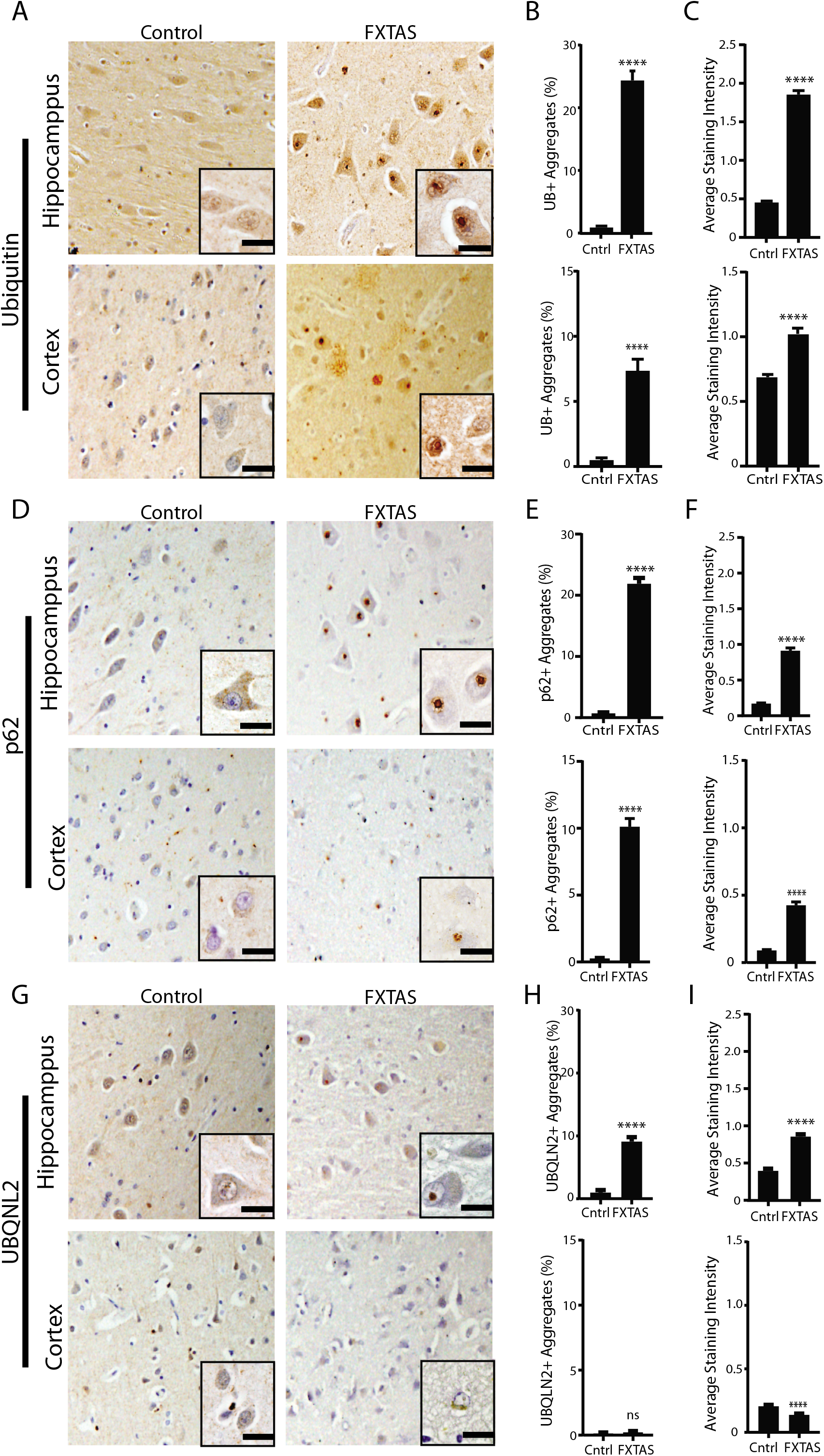
Staining for pathological markers Ubiquitin, p62, and UBQLN2 in FXTAS brain. A. Representative images from control (left) and FXTAS (right) hippocampus (upper panels) and cortex (lower panels) stained for ubiquitin positive aggregates. Nuclei stained with hematoxylin. Inset-60x magnification. Scale bar 20μm. B. Quantification of A represented as percent neurons with aggregates. Data from hippocampus (top) and cortex (bottom). Results expressed as means ± SEM; Student t-test **** p<0.0001 C. Graph showing average staining intensity for ubiquitin in hippocampus (top) and cortex (bottom). Results expressed as means ± SEM; Student t-test **** p<0.0001 D. Representative images from control (left) and FXTAS (right) hippocampus (upper panels) and cortex (lower panels) stained for p62 positive aggregates. Nuclei stained with hematoxylin. Inset-60x magnification. Scale bar 20μm. E. Quantification of D represented as percent neurons with aggregates. Data from hippocampus (top) and cortex (bottom). Results expressed as means ± SEM; Student t-test **** p<0.0001 F. Graph showing average staining intensity for p62 in hippocampus (top) and cortex (bottom). Results expressed as means ± SEM; Student t-test **** p<0.0001 G. Representative images from control (left) and FXTAS (right) hippocampus (upper panels) and cortex (lower panels) stained for UBQLN2 positive aggregates. Nuclei stained with hematoxylin. Inset-60x. Scale bar 20μm H. Quantification of G represented as percent neurons with aggregates. Data from hippocampus (top) and cortex (bottom). Results expressed as means ± SEM; Student t-test **** p<0.0001, ns-not significant I. Graph showing average staining intensity for Ubqln2 in hippocampus (top) and cortex (bottom). Results expressed as means ± SEM; Student t-test **** p<0.0001

To quantify the relative accumulation and intensity of staining for these three proteins in FXTAS, we performed rater blinded quantification of staining for each marker in both FXTAS and control samples from the hippocampus and cortex. A large fraction of FXTAS hippocampal neurons contained inclusions, with ubiquitin, p62, or UBQLN2 aggregates present in 25%, 20%, or 8%, of neurons (>300) counted in each of 4 FXTAS cases, respectively (Fig. 1B, 1E, and 1H). This was significantly higher compared to controls. Generally, a higher percentage of control neurons have no staining with any of these antibodies. The average staining intensity for each antibody was significantly higher in FXTAS tissue compared to control tissue (Fig. 1C, 1F, 1I). Results are summarized in Table 1.

**Table 1:**
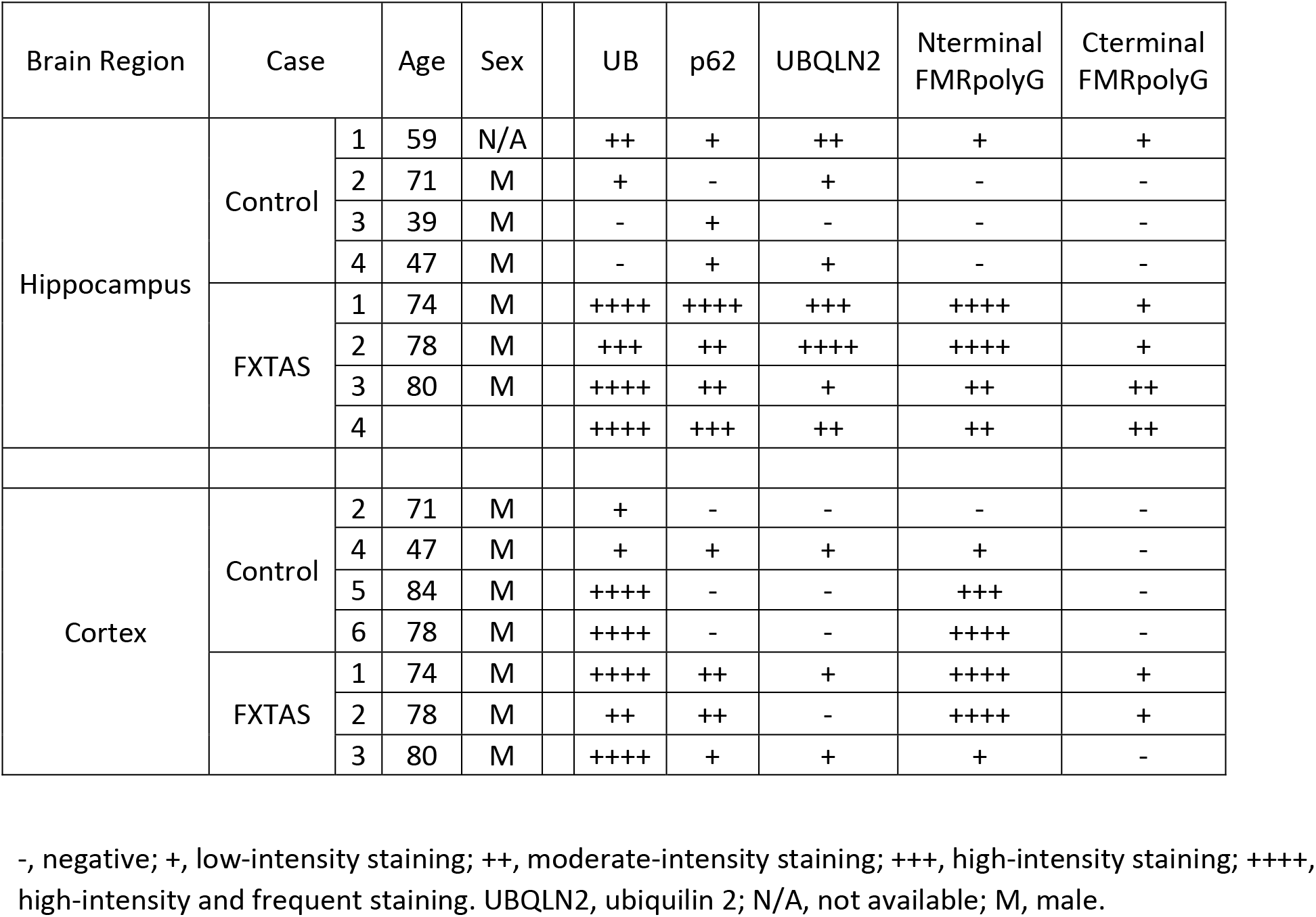
Summary of antibody staining in control and FXTAS patient tissue

To better characterize the role of RAN translation products in FXTAS, we generated a series of new polyclonal antibodies against FMRpolyG and FMRpolyA. Three previously published mouse monoclonal FMRpolyG antibodies were generated using epitopes just N-terminal (8FM) or C-terminal (9FM and 2J7) to the repeat element. Qualitative analysis in 2 FXTAS cases with 2J7 estimated that ~30% of the inclusions in FXTAS stained positive for FMRpolyG[29]. However, it was unclear whether the remaining inclusions were the result of alternative RAN translation proteins, repeat RNA triggered aggregates, or epitope masking that precluded recognition by the FMRpolyG antibody. Consistent with the latter concept, the N-terminal antibody had greater staining in FXTAS cases. We therefore used larger epitopes involving the N-terminal region (NTF1) and C-terminal region (CTF1) of FMRpolyG, under the assumption that some parts of the protein might be subject to proteolytic cleavage. The exact epitopes used are shown in Fig. 2A, underlined in red. A unique feature of the NTF1 and CTF1 was that each epitope also included a portion of the polyglycine repeat. Western blot analysis of HEK293 cell lysates expressing either EGFP-N1 or FMRpolyG fused to GFP showed both CTF1 and NTF1 antibodies specifically recognized FMRpolyG (Fig. 2B, 2D). NTF1 had some nonspecific staining compared to CTF1 by immunoblot. By immunofluorescence, both antibodies recognized cells expressing FMRpolyG (Fig. 2C, 2E).

**Figure 2.**
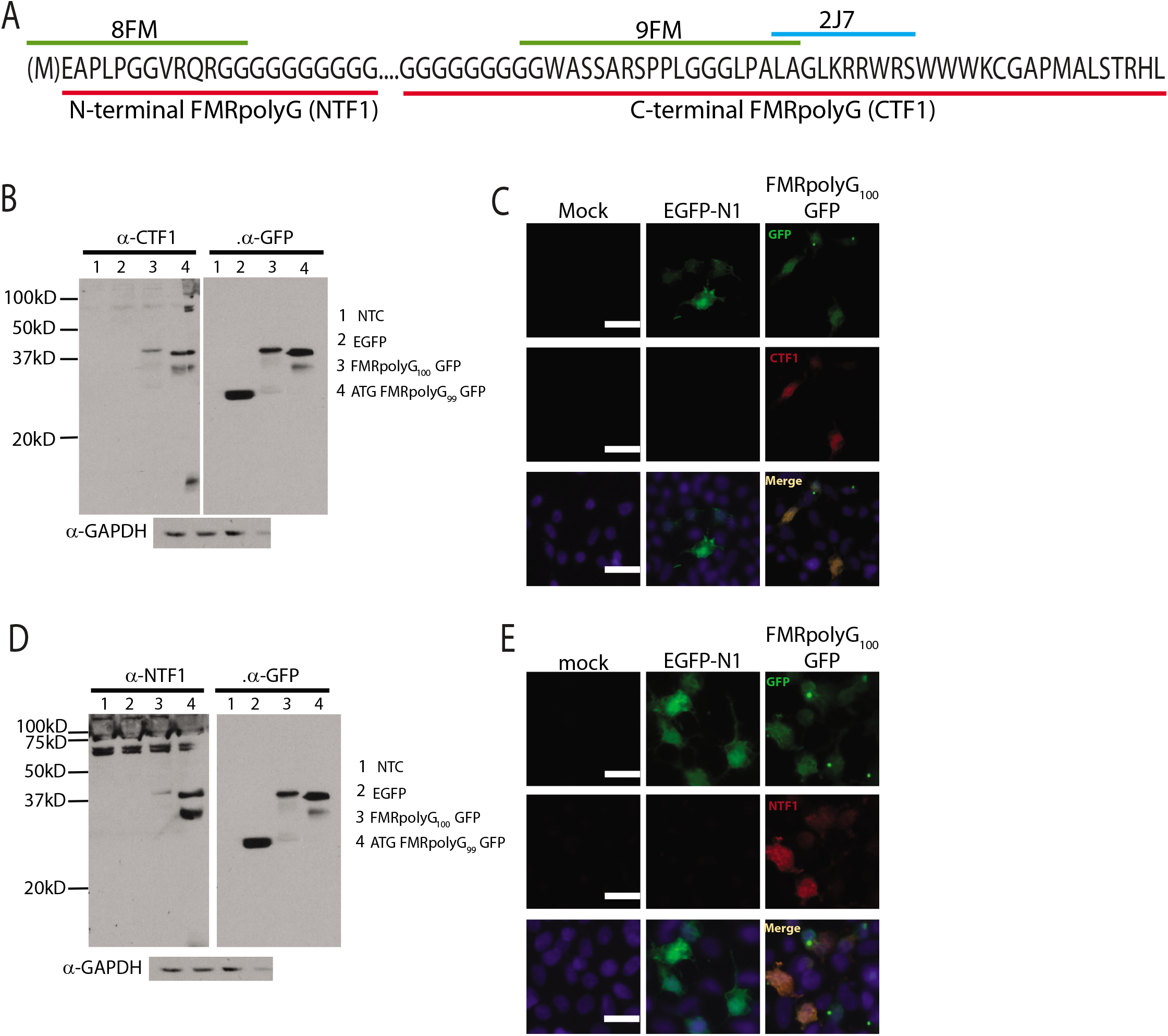
Generation and characterization of new FMRpolyG antibodies. A. Schematic showing epitopes for published (8FM, 9FM and 2J7) and novel (NTF1, N-terminal FMRpolyG and CTF1, C-terminal FMRpolyG) antibodies against FMRpolyG. Epitope sequence is underlined (green line - 8FM and 9FM, blue line-2J7, red line - N and C terminal FMRpolyG antibodies used in this paper). B. Western blot using CTF1 and GFP antibodies in HEK293 cells transfected with a nontemplate control (NTC-lanes1), EGFP-N1 plasmid (lanes 2), FMRpolyG 100GFP (lanes 3) and ATG FMRpolyG_99_GFP (lanes 4). GAPDH used as loading control. C. Immunocytochemistry of Mock, EGFP-N1 and FMRpolyG _100_GFP transfected HEK293 cells using CTF1 antibody. Nuclei stained using DAPI. Scale in C is 10μm. D. Western blot using NTF1 and GFP antibodies in HEK293 cells transfected with a nontemplate control (NTC-lanes1), EGFP-N1 plasmid (lanes 2), FMRpolyG _100_GFP (lanes 3) and ATG FMRpolyG_99_GFP (lanes 4). GAPDH used as loading control. E. Immunocytochemistry of Mock, EGFP-N1 and FMRpolyG _100_GFP transfected HEK293 cells using NTF1 antibody. Nuclei stained using DAPI. Scale in E is 10μm.

To validate the specificity of these antibodies, we performed a number of additional controls. A potential problem with polyclonal antibodies, in particular, is non-specific binding of the antibody to proteins not containing the antigen. The specificity can be determined by incubating the antibody with a peptide bearing the recognition sequence of the antibody. The antibody would bind the peptide and would not be able to bind the antigen in either protein lysates or human tissue. After incubating both NTF1 and CTF1 with their respective peptides, neither antibody recognized either FMRpolyG containing lysates (Fig. 3A, 3D) or FXTAS tissue (Fig 3B, 3E). Pre-immune sera for CTF had no staining in control or FXTAS hippocampal tissue (Fig. 3C) while pre-immune sera for NTF1 had light cytoplasmic staining in both control and FXTAS tissue (Fig. 3F).

**Figure 3.**
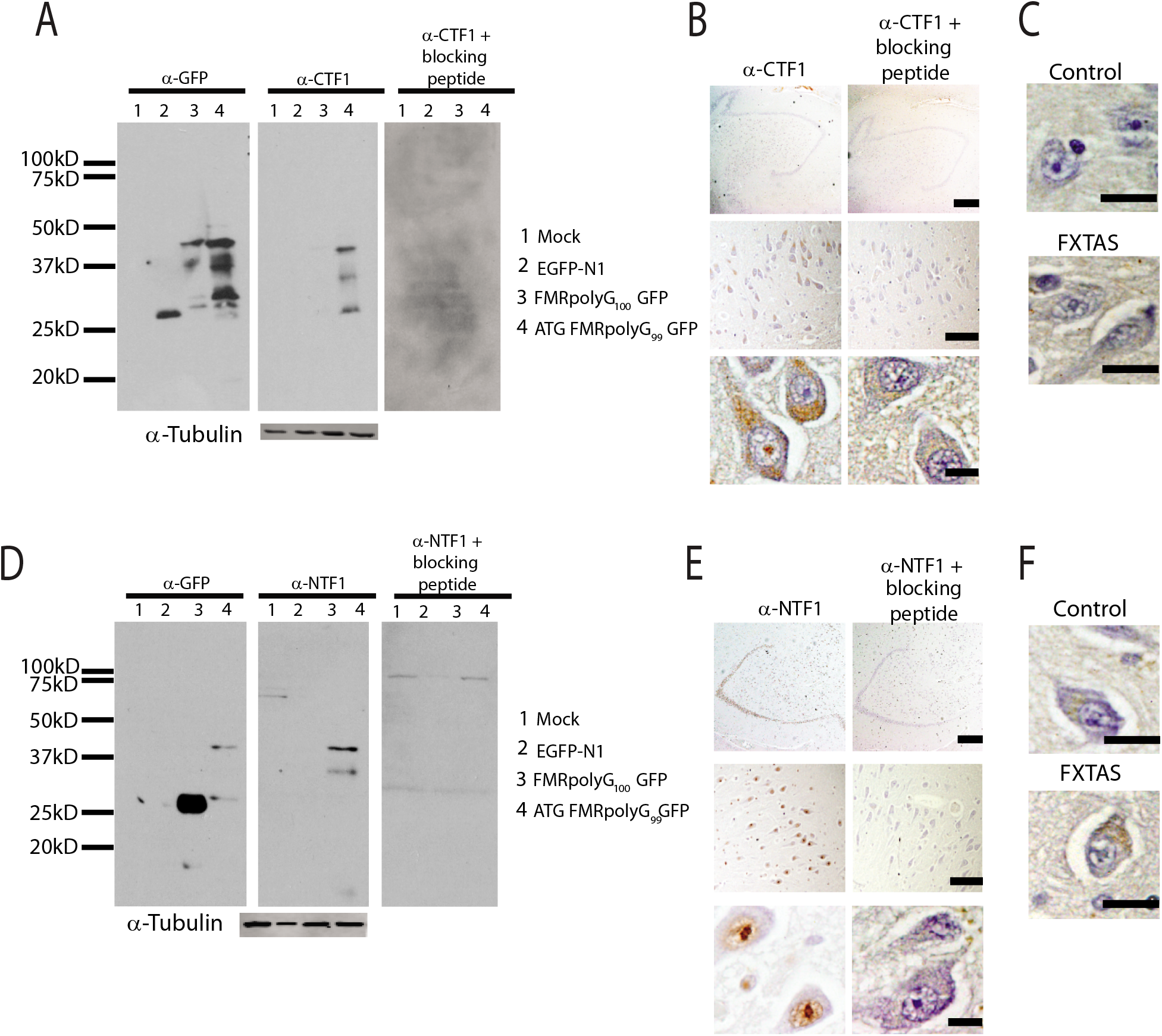
Validation of new FMRpolyG antibodies on FXTAS brain tissue. A. Western blot using GFP and CTF1 with and without blocking peptide in HEK293 cells that were either mock transfected (lanes1) or transfected with EGFP-N1 plasmid (lanes 2), FMRpolyG 100GFP (lanes 3) and ATG FMRpolyG_99_GFP (lanes 4). Tubulin used as loading control. B. Representative brain images from FXTAS patients stained using CTF1 antibody with and without blocking peptide at 4x (top), 20x (middle) and 60x (bottom) magnification. Nuclei stained with hematoxylin. Scale bars are 500μm, 100μm and 20μm respectively. C. Staining of control and FXTAS brain tissue using pre-bleed serum from CTF1 antibody production. Nuclei stained with hematoxylin. Scale bar-20μm D. Western blot using GFP and NTF1 with and without blocking peptide in HEK293 cells that were either mock transfected (lanes1) or transfected with EGFP-N1 plasmid (lanes 2), FMRpolyG _100_GFP (lanes 3) and ATG FMRpolyG_99_GFP (lanes 4). Tubulin used as loading control. E. Representative brain images from FXTAS patients stained using NTF1 antibody with and without blocking peptide at 4x (top), 20x (middle) and 60x (bottom) magnification. Nuclei stained with hematoxylin. Scale bars are 500μm, 100μm and 20μm respectively. F. Staining of control and FXTAS brain tissue using pre-bleed serum from NTF1 antibody production. Nuclei stained with hematoxylin. Scale bar 20μm.

With the newly validated antibodies, we looked to see if they could recognize FMRpolyG in patient FXTAS brain tissue. In hippocampal and cortical neurons, CTF1 exhibited significant cytoplasmic staining in both control and FXTAS tissue. However, CTF1 also detected FMRpolyG aggregates in FXTAS neurons (Fig. 4A, 4D). Overall, CTF1 positive inclusions were relatively rare, with approximately 3% and 1.2% of FXTAS neurons exhibiting CTF1 positive aggregates in the hippocampus and cortex, respectively, by blinded quantification (Fig. 4B, 4E). This was significantly greater than that observed in control tissues. The overall intensity of staining was significantly higher in FXTAS cortex and hippocampus compared to control (Fig. 4C, 4F).

**Figure 4.**
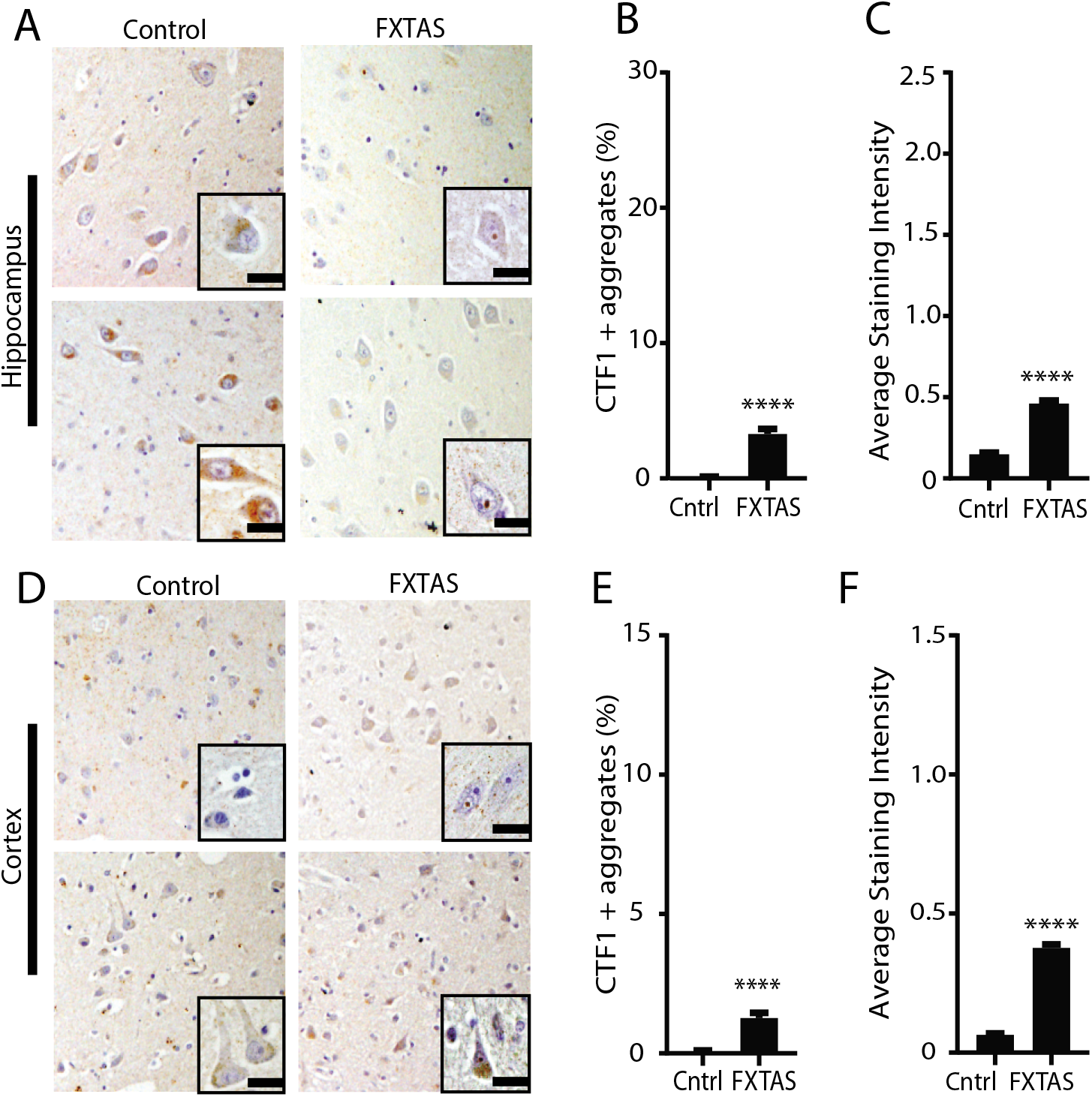
C-terminal targeted FMRpolyG antibody staining in FXTAS tissue. A. Representative images from control (left) and FXTAS (right) hippocampus stained with CTF1 antibody. Nuclei stained with hematoxylin. Inset-60x magnification. Scale bar 20μm. B. Quantification of A represented as percent neurons with CTF1 positive aggregates in hippocampus. Results expressed as means ± SEM; Student t-test **** p<0.0001 C. Graph showing average staining intensity for CTF1 positive aggregates in hippocampus. Results expressed as means ± SEM; Student t-test **** p<0.0001. D. Representative images from control (left) and FXTAS (right) cortex stained for CTF1 positive aggregates. Nuclei stained with hematoxylin. Inset-60x magnification. Scale bar 20μm. E. Quantification of D represented as percent neurons with aggregates in cortex. Results expressed as means ± SEM; Student t-test **** p<0.0001. F. Graph showing average staining intensity for CTF1 positive aggregates in cortex. Results expressed as means ± SEM; Student t-test **** p<0.0001.

In contrast to the CTF1 antibody, NTF1 staining of FMRpolyG was qualitatively different. Staining of hippocampal and cortical neurons with NTF1 showed low levels of cytoplasmic staining in control tissue. However, in some control cases, cortical neurons had moderate diffuse nuclear staining without aggregate formation. In FXTAS cases, NTF1 avidly stained intranuclear FMRpolyG positive aggregates in both the hippocampus and cortex (Fig. 5A, 5D). The percentage of neurons with FMRpolyG positive aggregates (17%) was similar to the percentages seen with ubiquitin (25%) and p62 (21%) and was significantly higher than control tissue (Fig. 5B) or staining with other published antibodies (data not shown). In the cortex, fewer total neurons had aggregates but the percentage of neurons with FMRpolyG aggregates in FXTAS cases was comparable to staining with Ubiquitin and p62 and much greater than control cases (Fig. 5E). The average staining intensity in FXTAS cortex and hippocampus was significantly higher than in controls by genotype and antibody blinded analysis (Fig. 5C, 5F).

**Figure 5.**
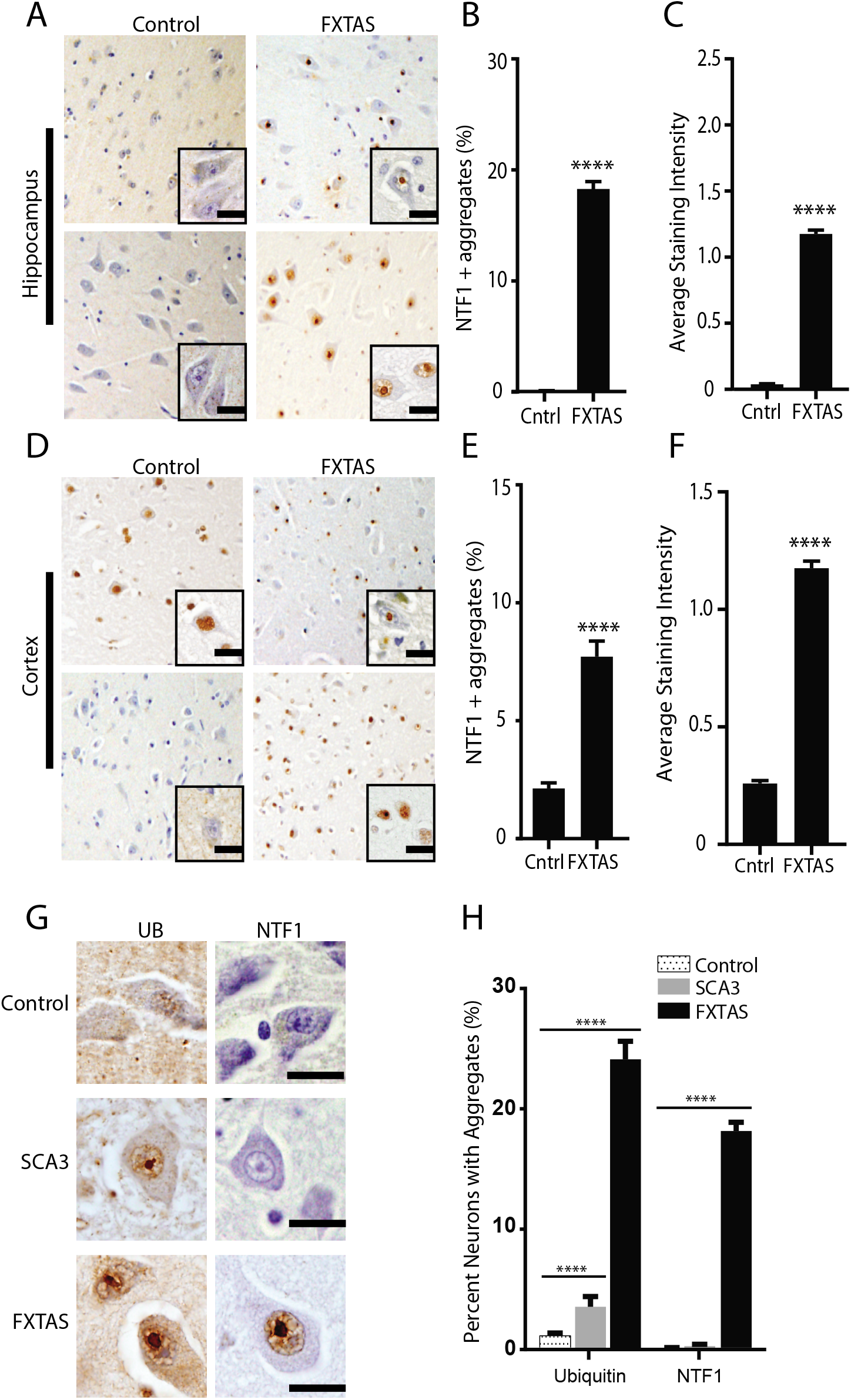
N-terminal targeted FMRpolyG antibody staining in FXTAS tissue. A. Representative images from control (left) and FXTAS (right) hippocampus stained with NTF1 antibody. Nuclei stained with hematoxylin. Inset-60x magnification. Scale bar 20μm. B. Quantification of A represented as percent neurons with NTF1 positive aggregates in hippocampus. Results expressed as means ± SEM; Student t-test **** p<0.0001 C. Graph showing average staining intensity for NTF1 positive aggregates in hippocampus. Results expressed as means ± SEM; Student t-test **** p<0.0001. D. Representative images from control (left) and FXTAS (right) cortex stained with NTF1 antibody. Nuclei stained with hematoxylin. Inset-60x magnification. Scale bar 20μm. E. Quantification of D represented as percent neurons with NTF1 positive aggregates in cortex. Results expressed as means ± SEM; Student t-test **** p<0.0001 F. Graph showing average staining intensity for NTF1 positive aggregates in cortex. Results expressed as means ± SEM; Student t-test **** p<0.0001. G. Representative images showing UB & NTF1 staining of control, SCA3 and FXTAS brain. Nuclei stained with Hematoxylin. Scale bars-20μm. H. Quantification of G showing percent neurons with UB and NTF1 positive aggregates in control, FXTAS and SCA3 tissues. Results expressed as means ± SEM; Student t-test **** p<0.0001.

Specificity of NTF1 for FMRpolyG was also evaluated by staining tissue from an SCA3 patient. The pons is a region of the brain known to have ubiquitinated aggregates in SCA3. When this region was stained with NTF1, nuclear aggregates were not observed in SCA3 patient tissue (Fig. 5G, 5H).

Neurodegeneration of the cerebellum is a common feature in FXTAS cases. We therefore used these antibodies on patient cerebellar tissue (Fig. 6) In FXTAS, ubiquitin, p62, and NTF1 stained aggregates throughout the granular layer of the cerebellum (Fig. 6B). Rare aggregates were seen in Purkinje cells with ubiquitin and NTF1, but these were less abundant than staining observed in the granule cell layer, a finding largely consistent with published studies.

**Figure 6.**
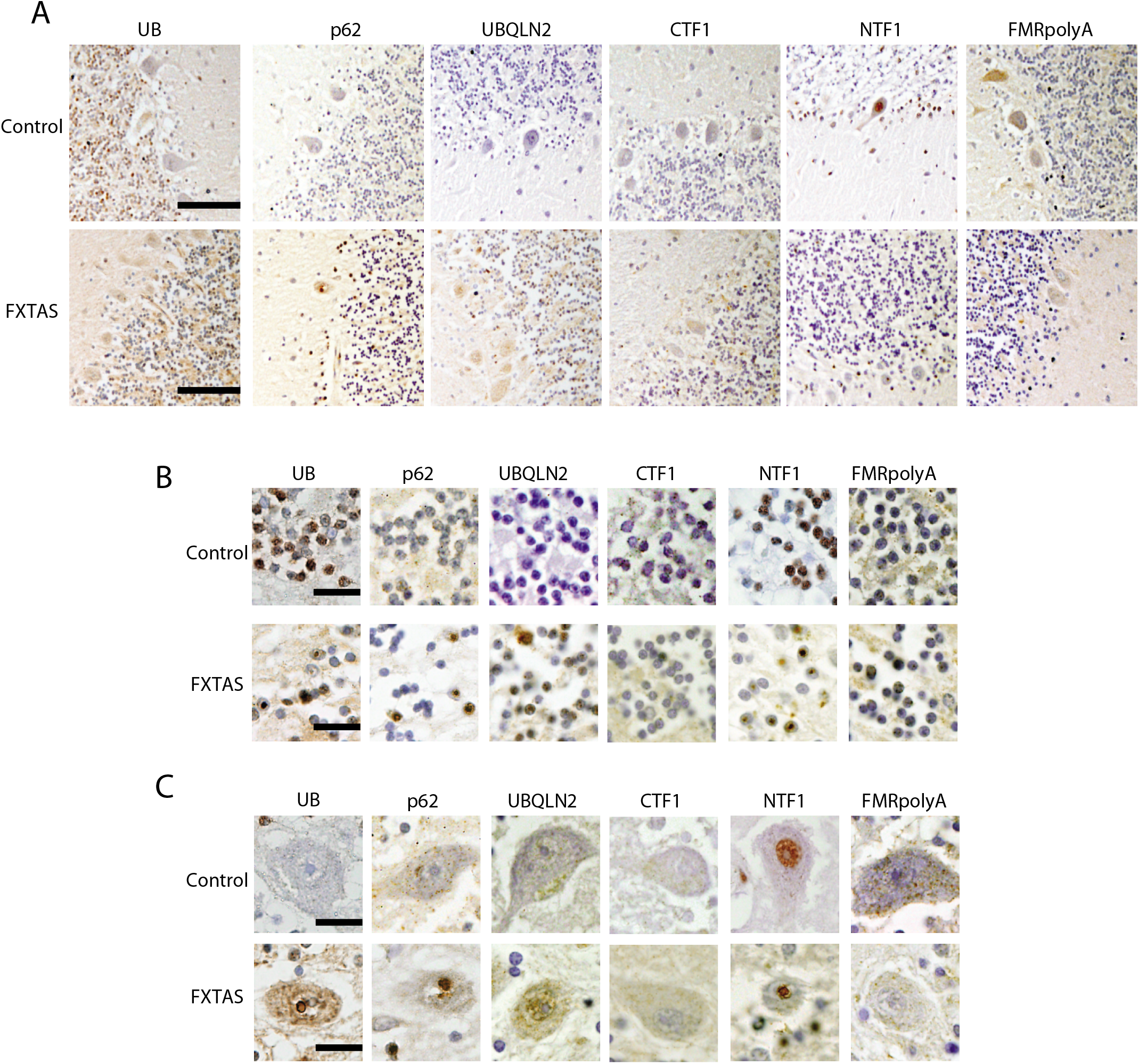
Cerebellar staining for FMRpolyG and other pathological markers. A. Representative 20x images of cerebellum stained with the indicated antibodies from control and FXTAS brain tissues. Scale bar is 100μm. B. Representative 60x images from the granular cell layer of control and FXTAS patients stained with the indicated antibodies. Scale bar is 20μm. C. Representative 60x images from the Purkinje cell layer of control and FXTAS patients stained with the indicated antibodies. Scale bar is 20μm.

Another protein generated by RAN translation is FMRpolyA. As with FMRpolyG, the contribution of FMRpolyA to FXTAS pathology was determined. A polyclonal antibody was generated against the C-terminal fragment of FMRpolyA (Fig. 7A). By western blot and immunocytochemistical analysis, the antibody specifically recognized FMRpolyA in transfected HEK293 cells (Fig. 7B, 7C). Incubating the antibody against FMRpolyA with the corresponding peptide prevented the antibody from recognizing FMRpolyA in HEK293 lysates or in human brain tissue (Fig. 7B, 7D). The pre-bleed serum has some light cytoplasmic staining in control and FXTAS patient tissue but aggregates were not seen (Fig. 7E).

**Figure 7.**
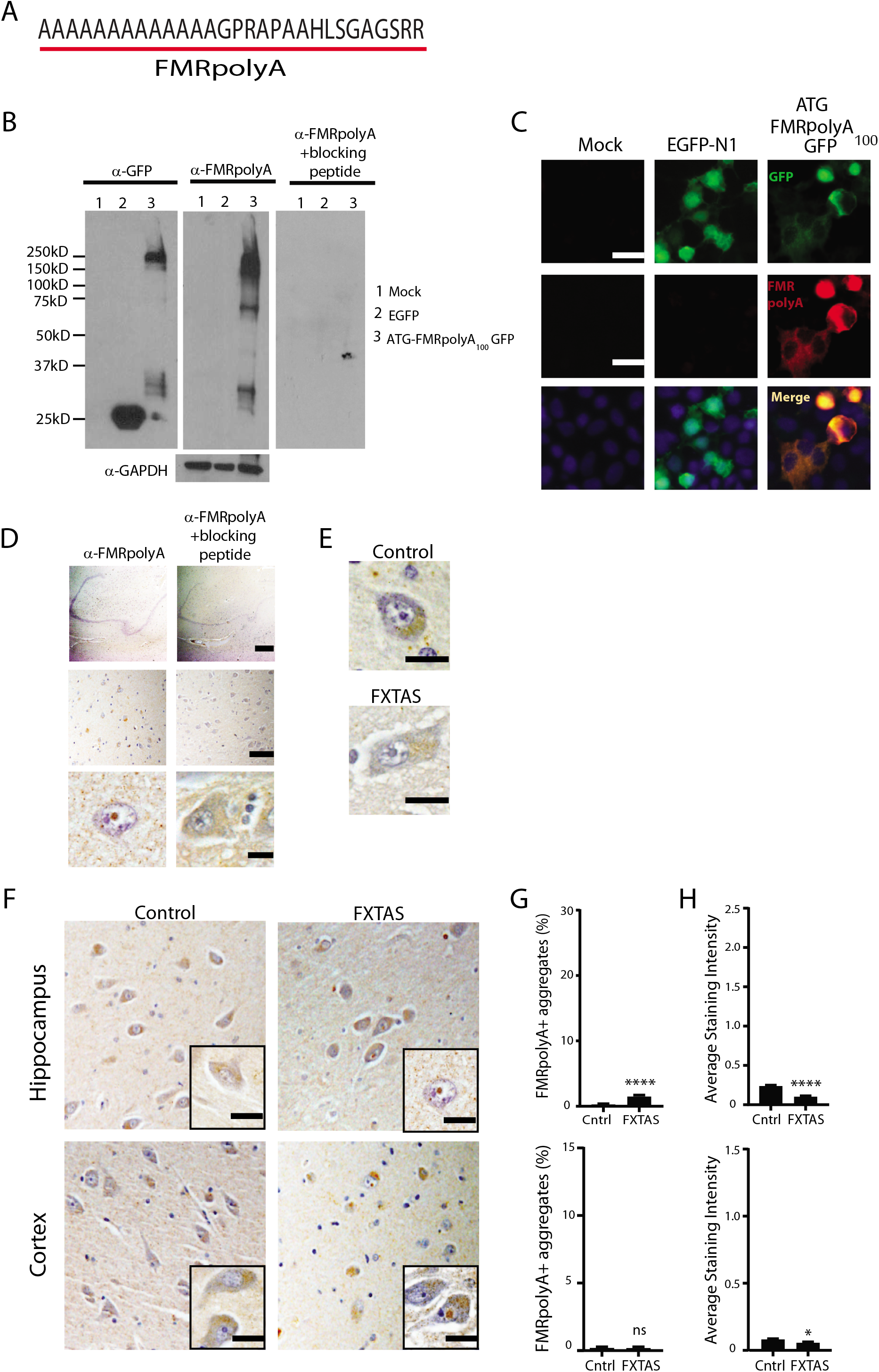
FMRpolyA staining in control and FXTAS tissues. A. Protein sequence of FMRpolyA epitope. B. Western blot using FMRpolyA antibody with and without blocking peptide and GFP in HEK293 cells that were either mock transfected (lanes1), transfected with EGFP plasmid (lanes 2) and ATG FMRpolyA_100_ GFP (lanes 3). GAPDH used as a loading control. C. Immunocytochemistry of Mock, EGFP-N1 and FMRpolyA _100_GFP transfected HEK cells using FMRpolyA antibody. Nuclei stained using DAPI. Scale bar is 10μm. D. Representative brain images from FXTAS patients stained using FMRpolyA antibody with and without blocking peptide at 4x (top), 20x (middle) and 60x (bottom) magnification. Nuclei stained with hematoxylin. Scale bars are 500μm, 100μm and 20μm respectively. E. Staining of control and FXTAS brain tissue using pre-bleed serum from FMRpolyA antibody production. Nuclei stained with hematoxylin. Scale bar 20μm. F. Representative images from control (left) and FXTAS (right) hippocampus (upper panels) and cortex (lower panels) stained with FMRpolyA antibody. Nuclei stained with hematoxylin. Inset-60x magnification. Scale bar 20μm. G. Quantification of F represented as percent neurons with aggregates. Data from hippocampus (top) and cortex (bottom). Results expressed as means ± SEM; Student t-test **** p<0.0001 H. Graph showing average staining intensity for FMRpolyA in hippocampus (top) and cortex (bottom). Student t-test * p< 0.05, ns not significant.

FMRpolyA exhibited light cytoplasmic staining in both control and FXTAS tissue. Intranuclear aggregates were very rare in hippocampal neurons (1.5%) and indistinguishable between control and FXTAS cortical neurons (Fig. 7F, 7G). The average staining intensity was actually decreased in FXTAS tissue compared to control tissue (Fig. 7H). FMRpolyA does not contribute significantly to FXTAS pathology.

To see if these antibodies are staining the same aggregates, co-immunofluorescence was performed. In FXTAS hippocampal neurons, ubiquitin positive aggregates were also positive for p62, UBQLN2, NTF1, CTF1, and FMRpolyA (Fig. 8A). Nearly all ubiquitin aggregates also stained positive for p62 and FMRpolyG with NTF1 whereas only 45% and 35% were positive for UBQLN2 or FMRpolyA, respectively (Fig. 8A). p62 co-localized to the same aggregates as NTF1 and UBQLN2 (Fig. 8B). Aggregates stained with NTF1 also were positive for ubiquitin, p62, and UBLQN2 (Fig. 8A, 8B). A summary of all staining observed is provided in Table 1.

**Figure 8.**
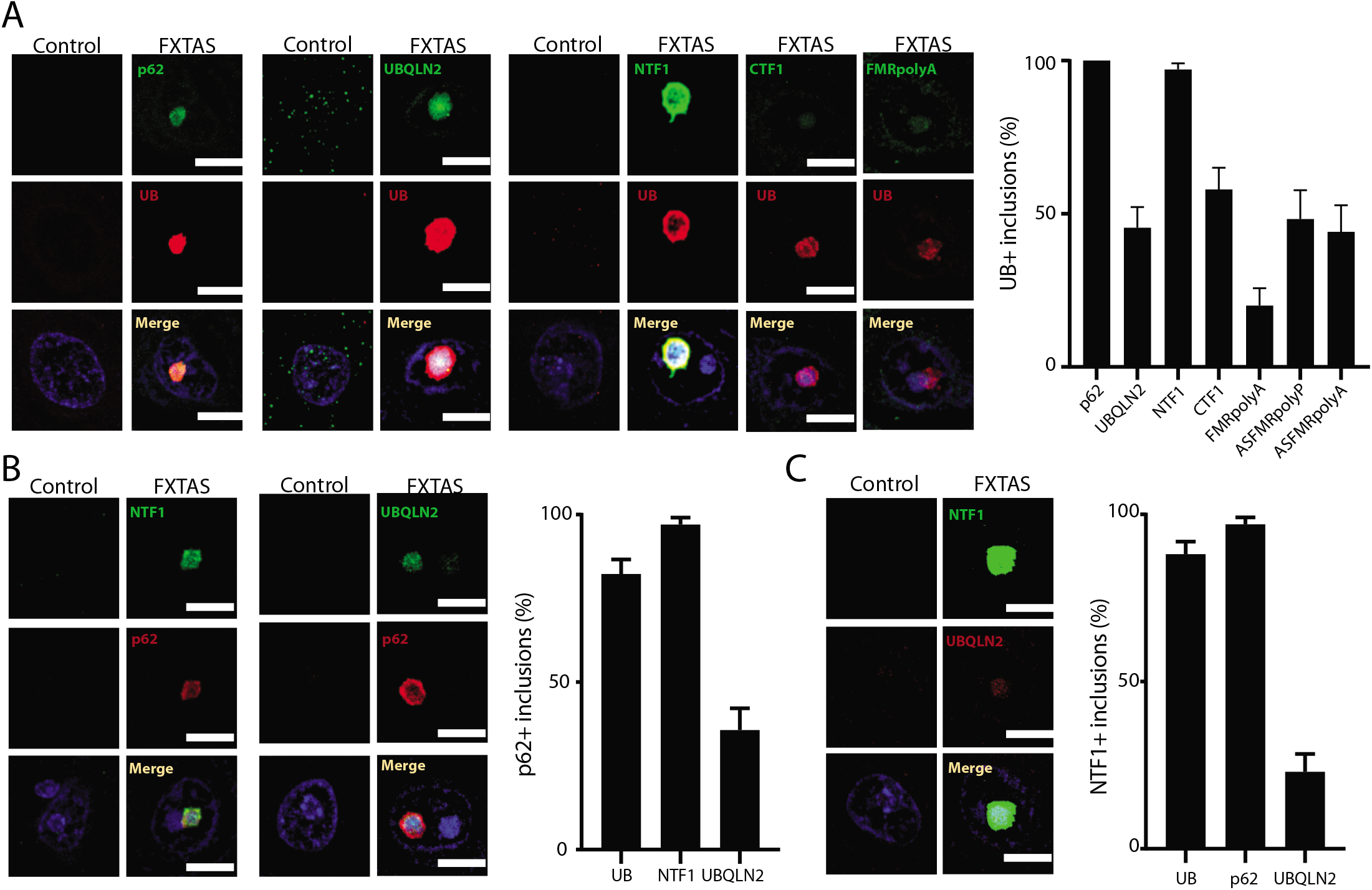
Co-localization of FMR antibodies. A. Immunofluorescence in control and FXTAS brain tissue co-stained with UB and either p62, UBQLN2, NTF1, CTF1 or FMRpolyA. Quantification (right) of each image representing the percent of UB positive aggregates that were also positive for each antibody. Nuclei were stained with DAPI. Scale bar = 10μm. B. Immunofluorescence in control and FXTAS brain tissue co-stained with p62 and either NTF1 or UBQLN2. Quantification (right) of image representing the percent of p62 positive aggregates that were also positive for NTF1 and UBQLN2 antibody. Nuclei were stained with DAPI. Scale bar = 10μm. C. Immunofluorescence in control and FXTAS brain tissue co-stained with NTF1 and UBQLN2. Quantification (right) of image representing the percent of NTF1 and UBQLN2 positive aggregates that were also positive for UB, p62 and UBQLN2. Nuclei were stained with DAPI. Scale bar = 10μm.

## Discussion

FXTAS is an under-recognized inherited neurodegenerative condition which lacks effective therapeutic options [8]. Accurately defining the pathological substrates for the disease in patients provides insight into how transcribed CGG repeat expansions elicit toxicity and drive neuronal death. Here we have evaluated a series of pathological markers predominantly in the hippocampus, cortex and cerebellum of FXTAS cases compared to controls in a rater-blinded fashion. These studies reveal that p62 and ubiquitin positive inclusions are present in a large percentage of neurons, with the greatest burden observed in pyramidal hippocampal neurons, consistent with past studies [4, 6, 7]. We find that FMRpolyG is detectable in the vast majority of these inclusions and in ~20% of all hippocampal neurons overall in FXTAS patients. In contrast, other RAN translation proteins generated from *FMR1* and *ASFMR1* mRNA are detectable in a much lower percent of inclusions, as is the ALS and autophagy-associated protein UBQLN2. In the context of published studies demonstrating that FMRpolyG is the most abundantly translated RAN protein from CGG repeats and that production of FMRpolyG is required for inclusion formation in *Drosophila*, transfected cells and both CGG KI and CGG transgenic mice, these data are consistent with a disease model whereby FMRpolyG production is a central contributor to ubiquitinated neuronal intranuclear inclusion formation in FXTAS.

We used two new polyclonal antibodies to FMRpolyG directed to the N-terminus (NTF1) and the C-terminus (CTF1) of the protein. Intriguingly, these two antibodies did not have the same patterns of staining. Specifically, NTF1 had diffuse nuclear staining in control cortex and hippocampus and robustly stained inclusions throughout the brain. In contrast, CTF1 had mild staining predominantly in the cytoplasm in control cases and stained inclusions in a lower fraction of neurons in FXTAS cases. The cause of this discrepancy is not yet known. Control studies using preimmune sera, pretreatment with a blocking epitope and immunoprecipitation of the protein followed by Mass Spectrometry (not shown) demonstrate that staining by both antibodies is specific to their epitopes. Moreover, NTF1 does not stain ubiquitinated inclusions in a different neurodegenerative disease, SCA3. In addition, both epitopes are unique within the proteome and even fragments of their epitopes do not clearly overlap with other nuclear or cytoplasmic proteins. These differences could reflect different affinities of the antibodies for their epitope targets. Arguing against this hypothesis is the finding that these staining patterns appear consistent with previously published antibodies, where monoclonal FMRpolyG C-terminal targeted antibodies have lower staining of inclusions and a diffuse cytoplasmic pattern in FXTAS tissues while the N-terminal targeted antibody more robustly stained inclusions. Access to the indicated epitopes in tissue may be different due to protein interactions or characteristics of the inclusions formed. Alternatively, FMRpolyG may undergo proteolytic cleavage in cells into smaller fragments that remove the C-terminal component of the protein and leave an N-terminal fragment consisting mostly of the polyglycine expansion fragment. Of note, this polyglycine component is critical for inclusion formation in overexpression systems [9, 21, 29].

When the 5’UTR of FMR1 is placed above GFP or nanoluciferase and expressed from plasmids or *in vitro* transcribed mRNA, FMRpolyG production occurs at both normal and expanded CGG repeat sizes in the absence of an AUG codon, suggesting that the protein could be made normally from *FMR1* mRNA. Consistent with this hypothesis, we observe that both NTF1 and CTF1 have some staining in control cortical and hippocampal neurons. If FMRpolyG is produced at low levels normally, what functions, if any, it might have are unknown. Previous work suggests that FMRpolyG interacts with a variety of proteins, including the nuclear lamin associated factor LAP2β. However, the interactome of this protein was defined in the setting of fusion of FMRpolyG to GFP with an expanded CGG repeat and with overexpression in HEK293 cells. As such, the relevance of these interactors to the native repeat size in human neurons or brain is not clear. Future studies will be needed to define what, if any, functions and interactions FMRpolyG has in neurons under normal contexts.

We also describe a new antibody raised against FMRpolyA. This protein is made less efficiently than FMRpolyG in RAN translation reporter assays and its production is much more dependent on an expanded CGG repeat [15]. Moreover, it is significantly less aggregation prone than FMRpolyG in GFP fusion studies in transfected cells [29]. Consistent with these features, we observe very little FMRpolyA in control or FXTAS brains and only a small percentage of inclusions stain positive for FMRpolyA. The staining observed is equivalent to or less than that observed even for RAN translation products generated from *ASFMR1* [16], whose transcript is produced at a significantly lower abundance that *FMR1* in both patients and controls. These findings are consistent with a prior study suggesting FMRpolyA is not readily observed in most FXTAS inclusions [21]. While these studies do not rule out a role for FMRpolyA in disease pathogenesis, they do suggest that it is not a central nucleator or major component of ubiquitinated inclusions in FXTAS.

This study has some limitations. Due to tissue access issues, we could not evaluate all of the antibodies described across all brain tissues and regions. In addition, the study used a relatively small cohort of cases and controls. As such, the generalizability of the findings to other FXTAS cases and premutation carriers may be limited. In addition, the ideal control tissue for these studies would have been brain samples from a fully methylated Fragile X Syndrome patient, as such a patient would presumably produce no *FMR1* or *ASFMR1* mRNA. However, such cases are rare and were not available to us. As such, a potential component of non-specific staining for FMRpolyG in control human brains in particular cannot be ruled out.

In summary, this study describes three new antibodies raised against epitopes from RAN translation products generated from CGG repeat expansions in *FMR1* mRNA. They confirm that FMRpolyG is a prominent component of ubiquitinated inclusions in FXTAS. These new antibodies will be a valuable resource to the research community and extend previous studies suggesting a role for RAN translation in the pathogenesis of FXTAS and other nucleotide repeat expansion disorders.

## Abbreviations

AR: Antigen retrieval
CTF1: C-terminal fragment 1 of FMRpolyG
FXTAS: Fragile X-associated Tremor/Ataxia Syndrome
GAPDH: Glyceraldehyde 3-phosphate dehydrogenase
GFP: Green fluorescent protein
KI: Knock-in
NIIs: Neuronal intranuclear inclusions
NGS: Normal goat serum
NTF1: N-terminal fragment 1 of FMRpolyG
PBS-MC: Phosphate buffered saline with magnesium and calcium chloride
PVDF: Polyvinylidene difluoride
RAN: Repeat associate non-AUG translation
SDS-PAGE: Sodium dodecyl sulfate polyacrylamide gel electrophoresis
UTR: Untranslationed region

## Acknowledgements

We thank the University of Michigan Brain Bank, the University of Florida Brain Bank, and the Colombia University Brain Bank for access to tissues. We thank Edgardo Rodriguez for providing access to tissues and early discussions on this manuscript. We thank the Paulson lab for access to microscopic equipment. We thank Michael Kearse and everyone in the Todd lab for many thoughtful conversations and their collective wisdom. This work was funded by grants from the VA BLRD (1I21BX001841 and 1I01BX003231), the NIH (R01NS099280 and R01NS086810), and the Michigan Alzheimer’s Disease Center and Protein Folding Disease Initiative to PKT. GS was supported by fellowships from FRAXA and the National Ataxia Foundation.

## Author Contributions

AK designed and performed experiments, analyzed and interpreted data, and wrote the manuscript with input from GS and YZ. GS and YZ designed/performed experiments and analyzed/interpreted data. BB performed experiments. PKT conceived the project, provided funding, designed experiments, interpreted data, and wrote the manuscript. All authors edited the manuscript.

